# Numerically enhanced adaptive optics-based 3D STED microscopy for deep-tissue super-resolved imaging

**DOI:** 10.1101/653394

**Authors:** Piotr Zdankowski, Maciej Trusiak, David McGloin, Jason R. Swedlow

## Abstract

In stimulated emission depletion (STED) nanoscopy, the major origin of decreased signal-to-noise ratio within images can be attributed to sample photobleaching and strong optical aberrations. This is due to STED utilising both a high power depletion laser (increasing risk of photodamage), while the depletion beam is very sensitive to sample-induced aberrations. Here we demonstrate a custom-built 3D STED microscope with automated aberration correction that is capable of 3D super-resolution imaging through thick, highly aberrating, tissue. We introduce and investigate image denoising by block-matching and collaborative filtering (BM3D) to numerically enhance fine object details otherwise mixed with noise. Numerical denoising provides an increase in the final effective resolution of the STED imaging of 31% using the well-established Fourier ring correlation metric. Experimental validation of the proposed method is achieved through super-resolved 3D imaging of axons in differentiated induced pluripotent stem cells growing under a 80µm thick layer of tissue with lateral and axial resolution of 256nm and 300nm, respectively.

## Introduction

Over the past two decades, super-resolution microscopy (SRM)^1,2^ has revolutionised the realm of fluorescent imaging. Imaging below the resolution limit is now commonplace using a range of widely available commercial and bespoke-built SRM systems. Fluorescence nanoscopy has proven to consistently provide resolution in the region of 20-50nm in all dimensions^3–11^, while some advanced techniques reported resolution around 1nm^12,13^. Unfortunately, increased resolution comes with a price – SRM systems are significantly more sensitive to optical aberrations and scattering, thus weakened signal leads to longer acquisition times and increased image noise. Many of the most impressive and ground-breaking results are acquired in the near-field region, very close to the coverslip for very thin slices^6,12–19^ or nanodiamonds^20–22^. The majority of the SRM solutions report only a lateral resolution below the diffraction limit and are not able to image samples thicker than 10µm^23^. For many systems, this implementation is good enough, but a push for imaging more complex biological samples in 3D drives the need for more sophisticated instruments. Among SRM techniques, stimulated emission depletion (STED) microscopy^24,25,26^ shows the greatest potential in imaging of thick specimens (>10µm), as it carries all the benefits (and downsides) of confocal microscopy.

Illuminating a fluorescent molecule with the appropriate wavelength will excite the molecule from its ground state to the excited state. In classical fluorescence microscopy the excited molecule emits via spontaneous emission. In STED this excited molecule can be instead forced to return to the ground state (depleted) using stimulated emission. This is achieved by illuminating it with a beam at the correct wavelength, and is termed the ‘depletion beam’. By appropriately shaping the depletion beam to have a null in the centre, and ensuring good spatial and temporal overlap between the excitation and depletion beams, the effective 2D point spread function (PSF) of the excitation beam can be dramatically reduced. Commonly this is done with a Laguerre-Gaussian (LG) beam^27^ allowing for a lateral spatial resolution improvement. In order to obtain axial super-resolution in the STED microscope, one needs to generate a 3D optical void in the depletion beam along the optical axis. This can be achieved through, for example, the use of a bottle-beam^28^ employing a π-step phase mask. This results in an increase of the axial resolution^29^ compared to 2D STED based on a LG beam. Another approach for obtaining a sub-diffraction axial resolution in STED microscopy combines STED with 4-Pi microscopy^7,30,31^.

Bypassing the diffraction limit when imaging thick samples remains a great challenge. Combining structured illumination microscopy (SIM) with 2-photon excitation enabled imaging of the microtubules in the eye of a living zebrafish embryo with 150nm lateral and 400nm axial resolution at a depth of 100 µm^32^. In single molecule localization microscopy, utilization of self-interference allowed for imaging at 50 µm depth with 100nm localisation precision^33^, while application of adaptive optics enabled imaging through 100 µm of tissue in the central nervous system of a Drosophila with a lateral localization precision of 143nm^34^. Lateral STED imaging through thick samples has been successfully demonstrated using Laguerre-Gaussian (LG) depletion beams^35–37^ as well as Bessel depletion beams for imaging fluorescent beads at depths as much as 155 µm^38^. Adaptive optics approaches have been implemented to image a rat heart at a depth of 36µm^37^ while a glycerol immersion objective lens with correction collar was used to facilitate imaging brain synapses at 120µm^39^. A combination of 2-photon microscopy with STED allowed for imaging at 30 µm inside a mouse brain^40^ and at depth of 50 µm in acute brain slices^41^. Imaging of thick samples with axial super-resolution in STED microscopy is more demanding and requires implementation of adaptive optics (AO) for aberration correction^42–44^. We have recently reported an AO 3D STED microscope demonstrating substantial improvements in axial resolution in single cell imaging^45^.

In this contribution we extend the achievable imaging depth in a highly scattering regime with an improved 3D STED system based on dual adaptive optics enabling imaging of axons in differentiated human induced pluripotent stem cells (hiPSCs) at a depth of 80µm using aberration correction. In addition, we investigate the issue of beam-intensity and scattering related signal-to-noise reduction in STED images. A numerical enhancement scheme for augmenting the STED signal-to-noise ratio is proposed.

Since STED microscopy is a purely optical method and does not require any image post-processing for obtaining resolution below the diffraction limit, image processing methods for noise removal and signal improvement have not yet received significant attention. Techniques such as deconvolution could be a possible solution to the retrieval of the object details otherwise covered by image noise. The PSF of the STED microscope is not constrained by its optical parameters, but mostly is defined by the fluorescent probe or the imaging conditions. For that reason, the PSF cannot be determined using sub-diffraction sized fluorescent beads but must be extracted directly from the acquired image. Deconvolution of STED images has been reported^46–48^, but requires the recorded image to have point-like structures for the determination of the PSF, which are not always apparent. Additionally, it is difficult to unambiguously state which structures can be treated as point emitters. Currently, neural network methods have shown that image resolution can be successfully increased in STED and localisation microscopy using deep learning, however those techniques require a very large number of ground truth images for the exhaustive network training^49–51^. Here we propose an approach for noise reduction in STED images and augmentation of the object-related structures making use of the block-matching and collaborative 3D filtering (BM3D) algorithm^52^. It does not require estimation of the image PSF nor extensive training of a neural network, in comparison to other currently employed image processing methods.

## Methods

### Dual-SLM 3D STED microscope setup

Our STED microscope schematic is presented in Fig. 1. It employs ps-pulsed lasers using 766nm for depletion (PicoQuant VisIR 765 STED) and 635nm for excitation (PicoQuant LDH-P-C 640B). The depletion beam is modulated using a liquid crystal on silicon spatial light modulator (SLM 1, Hamamatsu X-10468-02). A half-wave plate is used for the rotation of the polarisation of the depletion laser in order to ensure that the depletion laser polarisation orientation matches the orientation of the optical axis of the liquid crystal of SLM 1. SLM 1, in the depletion beam path, is used for both shaping the depletion beam and performing aberration correction. The hologram displayed on SLM 1 is imaged onto the back focal plane of the microscope objective MO (Nikon CFI Plan Apo Lambda 100X Oil NA 1.45) using relay optics.

**Fig. 1.**
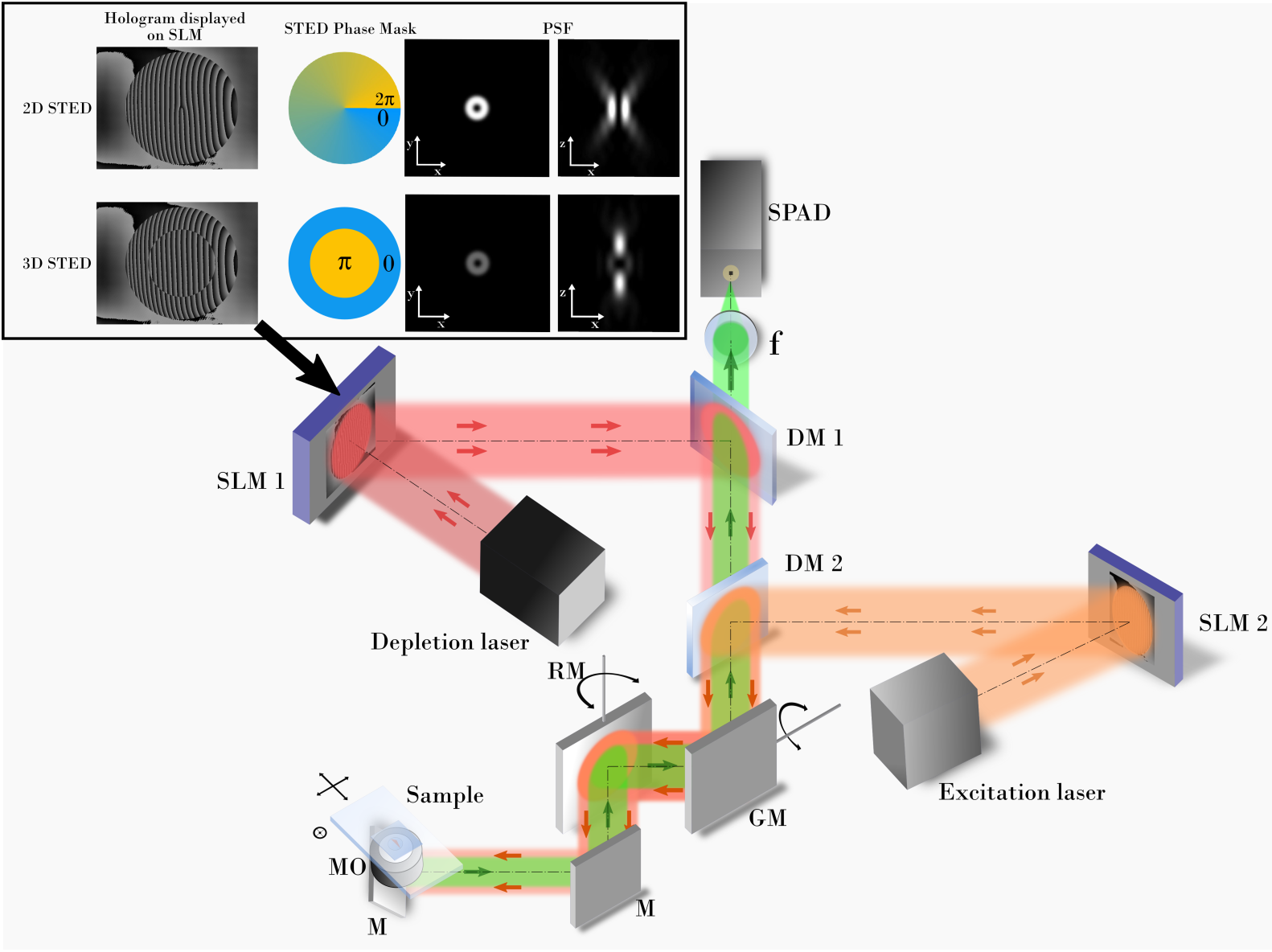
The optical setup of the custom built dual-SLM 3D STED microscope. M stands for steering mirrors, f – lens, MO – microscope objective, RM – resonant mirror, GM – pair of galvanometric mirrors, DM1-DM2 dichroic mirrors, SPAD – single photon avalanche diode, SLM 1 and SLM 2 – spatial light modulators. Detailed description of the setup and its elements can be found the supplementary materials. Inset show the typical depletion beam phase masks displayed on the SLM and the resulting beam shapes of the depletion beam.

After reflecting from SLM 1, the depletion beam reaches a short-pass dichroic mirror DM 1 (Semrock FF720-SDi01), which separates the fluorescence from the depletion beam. The STED beam passes through dichroic mirror DM 2 where it is combined with the excitation beam. The excitation laser is coupled into a single mode polarisation maintaining fibre and collimated at the output of the fibre. Then it is reflected from a second SLM (SLM 2, Boulder Nonlinear Systems 512×512), which is responsible for the aberration correction of the excitation beam. It is required for imaging thick specimens, as without a high-quality confocal image, super-resolution STED imaging will not be obtained. Dichroic mirror DM 2 (Chroma ZT647rdc-UF3) reflects the excitation beam into the main optical path. From the DM 2 plane, both excitation and depletion beams propagate coaxially. Two galvanometric mirrors (GM, ScanLab Dynaxis XS) and a 16kHz resonant mirror (RM, Electro-Optical Products Corp. SC-30) are used for the scanning of the beam across the sample. A quarter-wave plate is placed before the MO to convert the polarisation to circular at the BFP. Both beams then pass through the microscope objective, which is placed on a *z*-piezo stage (Piezoconcept HS1.70) used for focusing, into the sample. The sample is mounted on a motorised XY-stage. The fluorescence signal that originates at the sample plane is focused with the imaging lens *f* into a multi-mode fibre (MMF), which is used as a confocal pinhole. The other end of the MMF is attached to a single photon avalanche diode SPAD detector (Excelitas SPCM-780-13-FC).

### Adaptive optics-based aberration correction procedure

Imaging thick samples introduces aberrations that increase with sample thickness. This is mainly due to the local variations of the refractive index within the imaged sample. This problem becomes very significant in 3D STED microscopy, as the bottle-beam is especially sensitive to aberrations^53–56^. To correct aberrations in our microscope, we use a correction routine based on the ‘modal wavefront sensing’ method^57^. In this approach, the wavefront is described as a summation of orthogonal modes, e.g. Zernike polynomials. The SLM can modulate the wavefront according to the applied modes and correct for the aberrations that originate both in the optical system and the imaged sample^58–61^. The aberration correction protocol implemented in our microscope is based on previous work described by Gould *et. al.*^42^, Patton *et. al.*^44^ and ourselves^45^. Zernike polynomials are well-established modes that accurately describe optical aberrations. We correct the following aberrations: 1^st^ order astigmatism, coma and trefoil, 1^st^, 2^nd^ and 3^rd^ order spherical. Each of the Zernike polynomials is corrected one at a time using the square of the sum of total intensity of the image as a quality metric. Correction of the excitation beam is straightforward using the maximum of the quality metric. Correcting the depletion beam requires two steps. Firstly, before imaging of the sample, a coarse correction and alignment is performed using gold beads and the quality metric is maximised as in the case of the excitation beam. To correct the specimen induced aberrations it is imaged with both excitation and depletion beam turned on, but the depletion beam has no phase mask applied, i.e. it has a Gaussian profile. Then, instead of maximising the quality metric, we find its minimum. This way, the depletion efficiency is maximised and the aberrations are corrected. When performing the aberration correction routine, both excitation and depletion beam intensities are set to the minimum value which still enables the correction to be recorded, in order to minimise optical sample exposure and decrease the risk of possible photo-damage.

### Image acquisition and processing

Microscope control and image acquisition were carried out using custom software written in Labview. Raw images with the experiment metadata were stored in the OME-TIFF format^62^. OMERO software^63^ was used as an image library. Using the OMERO Matlab toolbox, the images were directly accessible in Matlab, which was used for the BM3D algorithm implementation and all image processing. We used the available implementation that was shared by its authors^52^.

## Results

### Noise problem and resolution quantification in STED imaging

The most common metric used for measuring the image resolution in STED microscopy has been the full width at half maxima (FWHM) calculation of a Gaussian or Lorentzian fit to the intensity cross-section of fine (nanometric sized), point-like and isolated object structures. While there are tools to measure the FWHM in situ without using fluorescent beads^64^, a problem appears when the imaged structure does not contain such fine components. Fluorescent beads can be used as a test object, though beads are usually much brighter in comparison to biological samples. What is more, bead images are acquired with much superior signal-to-noise ratio (SNR), and noise is not as big of a factor when calculating the resolution. For that reason, a far more comprehensive resolution evaluation metric, called Fourier ring correlation (FRC), has been proposed by Nieuwenhuizen *et al*.^65^ Originally, this metric was applied to localization based nanoscopy with Tortarolo *et al.*^66^ proposing FRC as a tool for the evaluation of the lateral resolution of STED images. FRC has also recently been proposed as a frequency domain filtering and as a helpful metric in the blind deconvolution of STED images.^48^ Comprehensive analysis proved it to be far superior than the FWHM calculation, which is why the FRC is used in this work to quantify the resolution. We make use of the Matlab tool that was developed and disseminated with the original FRC publication^65^.

Calculation of the FRC requires two identical images recorded independently of each other to correlate two image noise instances. Thanks to the nature of how images are acquired when scanning the sample using the resonant mirror, each raw image contains a mirror image^67^. Having independently acquired these twin images, they then need to be translated so that there is no drift between them, as it can influence the FRC result^66^. We use the subpixel registration algorithm based on the cross correlation of the images^68^. In this way we can be sure that drift does not influence the FRC calculation. Using the FRC as an effective resolution metric, one needs to define the criterion for an effective cut-off frequency that will correspond to the highest spatial frequency passed by the imaging system. All other frequencies below the set threshold will be treated as unresolvable due to their amplitude being too low. In the works that proposed using the FRC as a resolution metric in SRM, a fixed threshold of 1/7 of correlation value is reported to produce reliable results and is already well-established^48,65,66^.

When acquiring STED images, the resolution can be increased by increasing the intensity of the depletion laser. This argument, however, does not take into account the increased photobleaching and noise that arises as a consequence, which can significantly degrade the final resolution of the acquired image. Figure 2 presents the FRC analysis of the fluorescent bead images obtained at increasing depletion laser intensities.

**Fig. 2.**
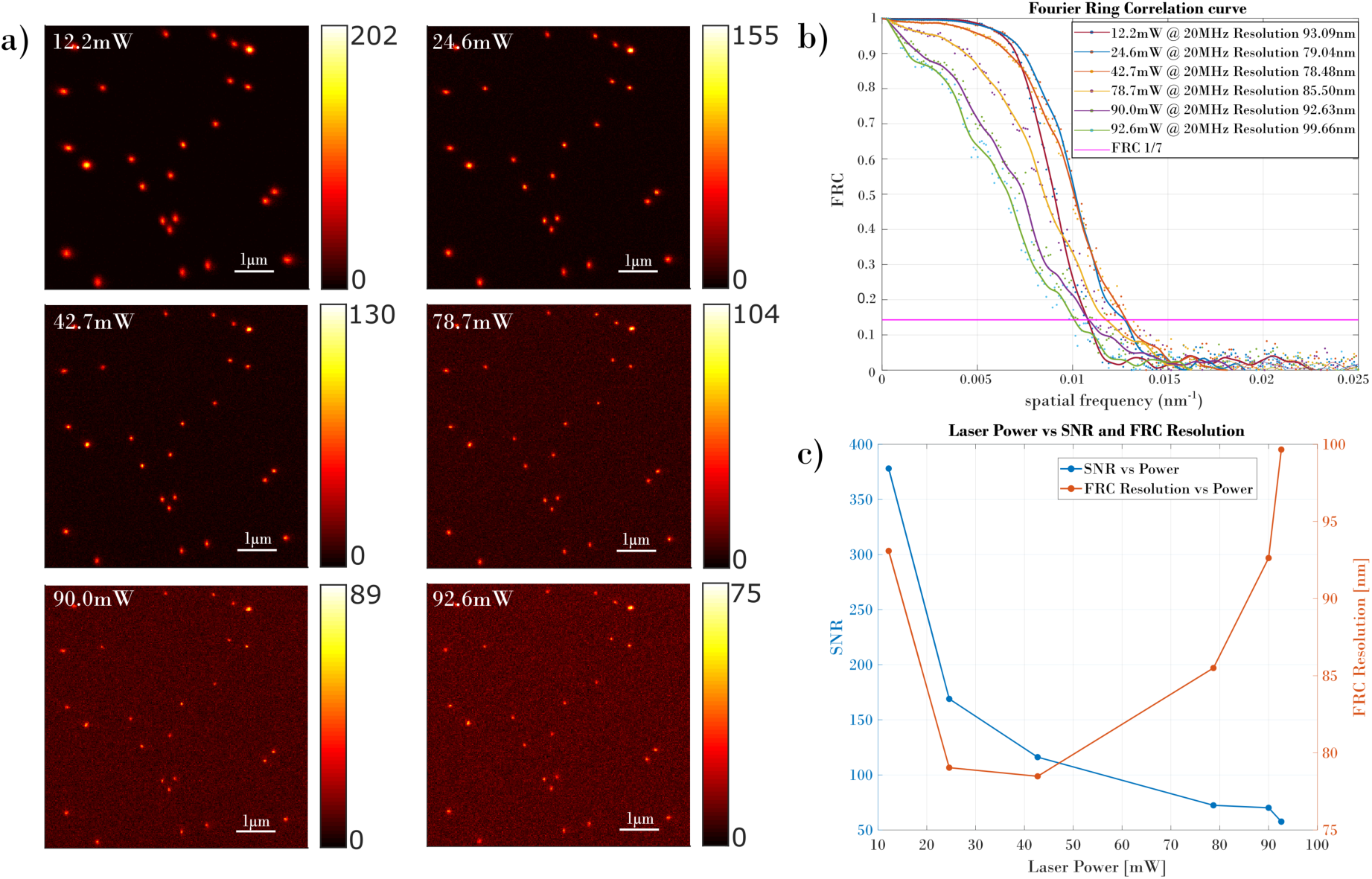
(a) Bead images acquired for different laser beam power (as shown in each image). Graph in (b) shows the microscope performance and effect of laser power related noise on the STED resolution determined by the FRC. Plots in (c) present the relation of depletion laser power and SNR (blue plot) and the relation between depletion laser power and FRC Resolution (red plot).

### Deep tissue imaging with dual-SLM 3D STED microscope

In order to visualise the impact that AO has on 3D STED imaging we imaged hiPSCs subjected to a differentiation protocol that produces dopaminergic (DA) neurons at high efficiency^69^. This procedure drives hIPSCs through a defined differentiation pattern, first producing neural precursors and then, by 30-35 days in culture, producing cells that express Tuj11 and tyrosine hydroxylase, a dopaminergic neuron marker. The procedure initiates by concentrating the cells into embryoid bodies (“EBs”), pseudo-tissues that are several cell layers thick. During the differentiation protocol, cells initially spread out from the EBs and then proliferate and differentiate, forming increasingly thick tissue. By ∼35 days in culture, the tissue is 50-80 um thick with differentiated neurons embedded in the tissue. Preparations of tissue grown on #1.5 coverslips were immunostained with Tuj-1 primary antibody, a neuron-specific tubulin marker, and with Aberrior STAR 635P^70^ as a secondary antibody and imaged in the 3D AO microscope. Fig. 3 shows the results obtained for imaging the sample with the confocal and 3D STED modes of microscope (employing aberration correction protocol and without it).

**Fig. 3.**
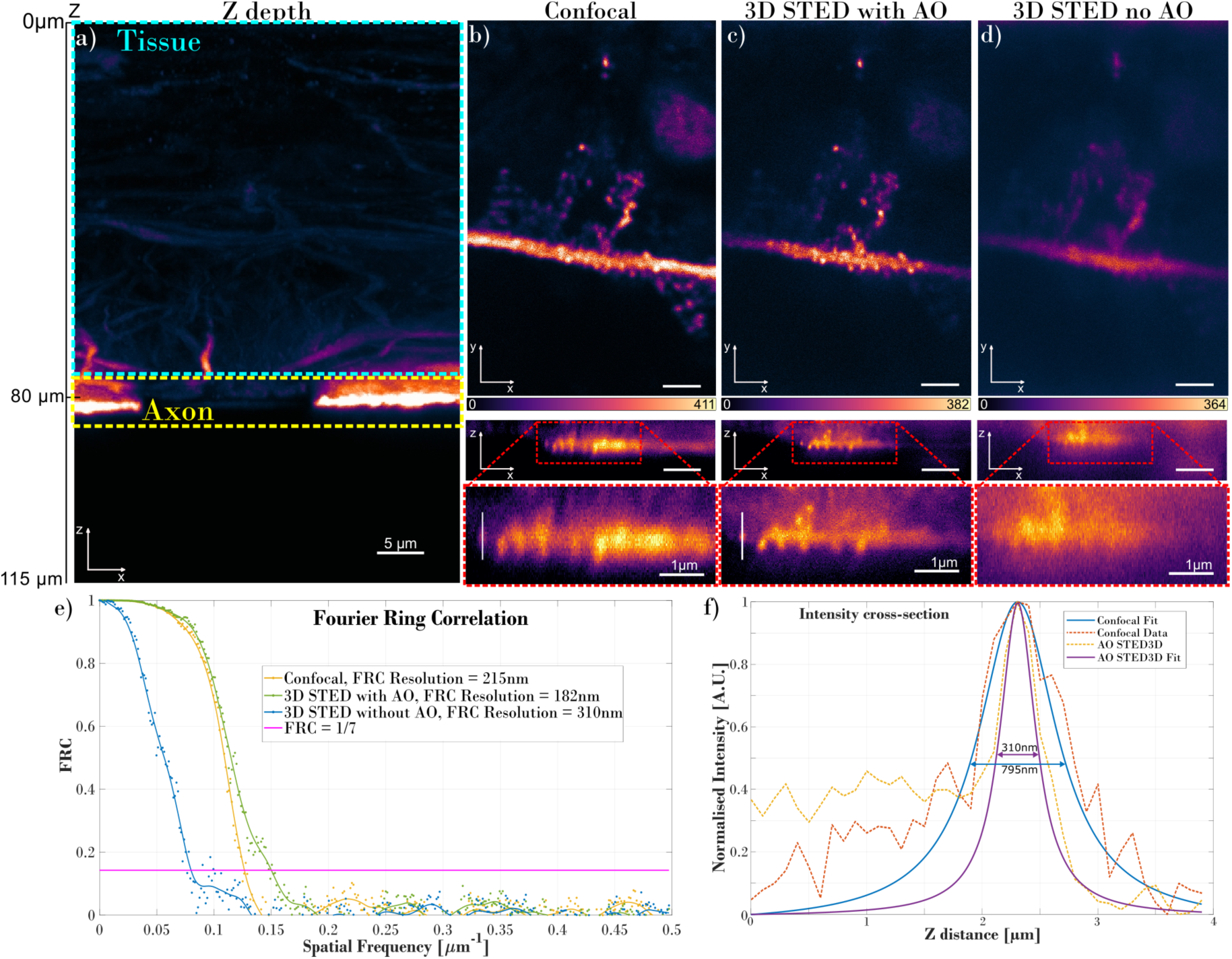
Comparison of the microscope performance when imaging through the aberrating specimen. (a) shows the structure of interest – an axon that grew above the layer of 80µm of tissue – in the XZ view. Top images in (b), (c) and (d) show the XY slice of the axon (top), while the XZ horizontal view is shown in the lower images. The dashed red rectangle marks the area that is zoomed in the lower XZ images. Plots in graph (e) present the calculated FRC for (b), (c) and (d); plots in graph (f) show intensity cross-sections and the Lorentzian fits used for the calculation of the FWHM. The position of the intensity cross-section is marked with the white vertical line in (b) and (c). Scalebar has 2µm size, unless stated otherwise.

The *xz* stack in Fig. 3(a) shows that the hiPSC4 cells grew an 80µm layer of tissue, above which it grew an axon. Figure 3(b) presents the confocal mode imaging outcome. As can be seen in the comparative image in Fig.3(c), AO enabled high quality and high-resolution imaging of the axon structure through the strongly aberrating 80µm layer of tissue. With no aberration correction the resulting image was overwhelmed by scattering-driven noise and aberration-driven blur, Fig. 3(d). The resolution in the z-direction can clearly be seen to have increased in Fig. 3(c) when comparing the confocal (Fig. 3(b)) and 3D STED images without aberration correction (Fig. 3(d)). The FRC graph shown in Fig. 3(e) compares the imaging FRC resolution calculated for each technique used. Confocal offers an FRC resolution of 215nm, uncorrected 3D STED gives 310nm, while AO 3D STED provides 182nm for the tissue slice shown in Fig. 3. The average FRC resolution over the whole volume of imaged axon (from z=80µm to z=83.5µm) reported 225nm, 323nm and 204nm for confocal, uncorrected and corrected 3D STED, respectively. Measuring the FWHM of the image feature in the z-direction, as shown in Fig. 3(f), allowed us to estimate the axial resolution as 795nm and 310nm for confocal and AO 3D STED images, respectively. The axial resolution had to be estimated by fitting the Lorentzian and calculating the FWHM, as the FRC metric can only be used for measuring the lateral resolution with equal pixel size in all directions. However, the FWHM establishes a good metric for the comparison of the performance of 3D confocal and AO 3D STED images. According to the FWHM measurement, the axial performance of the microscope using the AO 3D STED phase mask is 2.5 times better than when imaging in the confocal mode, but with AO correction. In the absence of AO correction resolution in the image plane and along the optical axis is severely degraded (Fig. 3(d)).

Figure 4 shows the amplitudes of Zernike coefficients for the depletion beam. Amplitudes shown in Fig. 4 (a) present the values used to correct optical system aberrations, disregarding the sample introduced aberrations, while Fig. 4 (b) also corrects the aberrations that were introduced by the imaged specimen. In order to obtain high quality and high-resolution images, strong spherical aberration introduced by the sample had to be compensated in our custom dual SLM system.

**Fig. 4.**
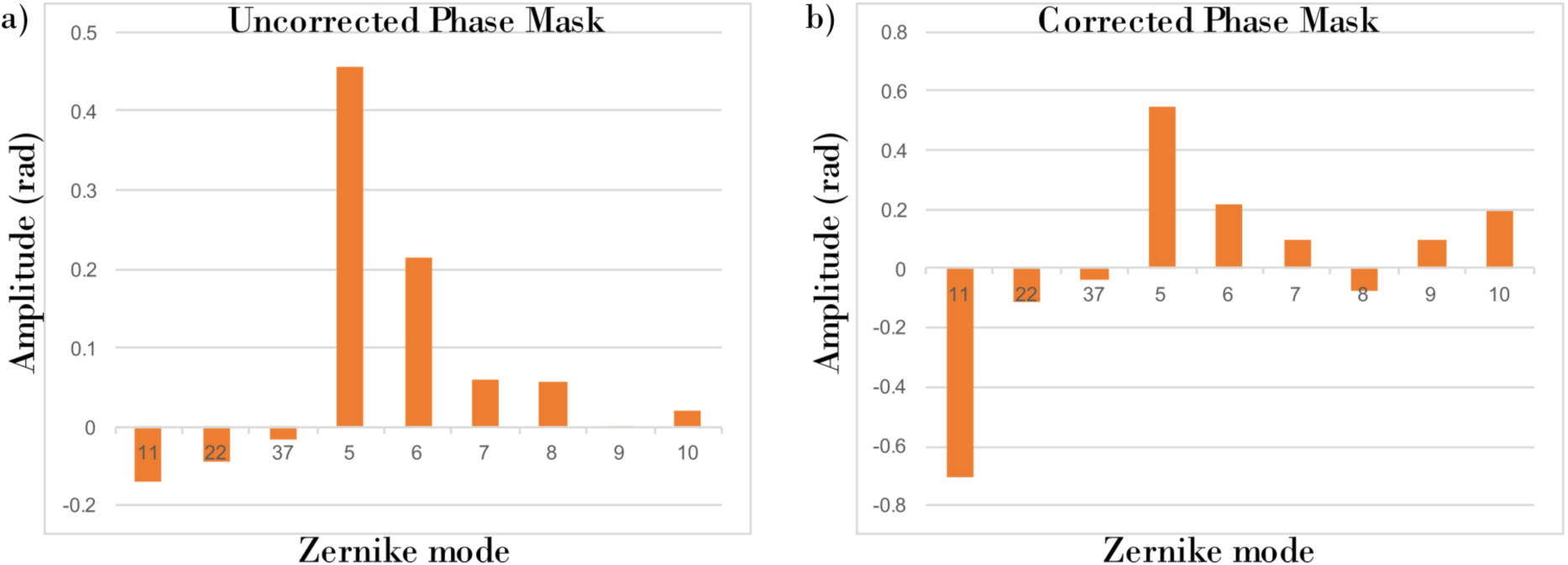
Zernike coefficients that were applied to the depletion beam SLM for uncorrected (a) and corrected (b) aberrations introduced by the imaged specimen, respectively.

Noise is significant when imaging through aberrating and optically dense structures and can suppress the spatial frequencies within the object structure, which will result in a decrease of the image resolution. Hence image denoising could be very beneficial to the resulting image and might enable retrieval of spatial frequencies that are otherwise mixed with the high level of noise.

### Image denoising using block-matching and collaborative filtering (BM3D)

The BM3D is a state-of-the-art algorithm widely used for denoising images^52^. It has been successfully used in speckle filtering in holography and holographic microscopy^71,72^, as well as in 2-photon microscopy^73^. The BM3D algorithm is performed in two steps. In the first step, the image is analysed for regions with local and non-local similarities called ‘blocks’ which can then be sorted into 3D arrays. These arrays then undergo a 3D linear integral transform followed by hard thresholding. Then the inverse 3D integral transform is carried out resulting in a first estimate of the ground-truth image. In the second step, the estimated image is put through a similar process with Wiener filtering replacing the hard thresholding. This time, the inverse transform gives the final denoised image exhibiting detail-preservation feature. Fig. 5 shows the graphical schematic of the operation of this 3D denoising algorithm. We anticipate that the BM3D filtering can be successfully used in the filtering of biological structures, since the cells tend to grow in a similar manner and have similar shape and morphological features which significantly simplifies the block matching procedure for 3D stack generation. Using a non-local search of similar image regions to the one under denoising, a crucial increase in differentiation between object and uncorrelated noise components in the integral transform coefficient domain is obtained. Here we use the basic, unmodified version of the BM3D algorithm, with σ (an estimate of the variance of the image noise) being the only variable that we want to modify. The algorithm was implemented in Matlab and disseminated upon original publication by the authors^52^.

**Fig. 5.**
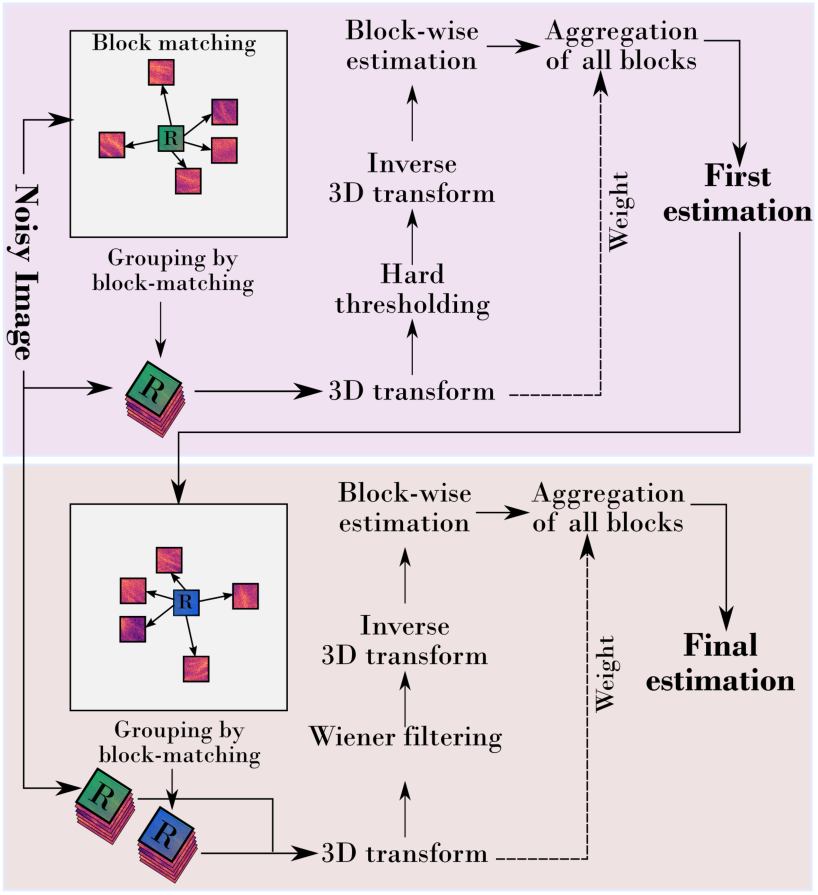
BM3D denoising algorithm schematic.

For evaluating the effectiveness of the BM3D denoising algorithm, we use the FRC metric for measuring the effective FRC resolution. Hence FRC can be also employed as a SNR improvement metric showing the increase of the effective resolution in the raw and denoised STED images.

In Fig. 6 we simulated white Gaussian noise and added it to the image in order to investigate the performance of the BM3D algorithm with the ground truth model enabling quantitative evaluation. A 2D STED image of the RPE-1 mitotic cell was acquired and then synthetically spoiled with controllable noise. The RPE-1 cells’ tubulin was stained with the Abberior STAR 635P secondary antibody. The Gaussian noise was selected as it is generated in a straightforward manner, has easily controllable statistics and efficiently damaged the image SNR. Random noise with the variance N (from N = 0 to N = 0.55) was simulated using Matlab with the *randn* function and then added to the image. Thanks to the noise simulation, we were able to evaluate the effectiveness of the BM3D denoising algorithm used on the STED images. Fig. 6 shows the impact of the noise on the effective image FRC resolution and the effectiveness of the BM3D denoising algorithm. Fig. 6(a) and 6(i) show the image without and with added noise with a variance of 0.25. Fig. 6(b) – (d) show the results of the denoising employing the BM3D algorithm with σ=5, σ=10 and σ=20, respectively, of the noise-free image (Fig. 6(a)). Fig. 6(j) – (l) demonstrate the BM3D denoising with σ=5, σ=10 and σ=20, respectively, of an image with added noise of 0.25 variance (Fig. 6 (i)). The yellow lines indicate the position of the intensity cross-sections shown in Fig. 6(e)-(h) and Fig. 6(m)-(p).

**Fig. 6.**
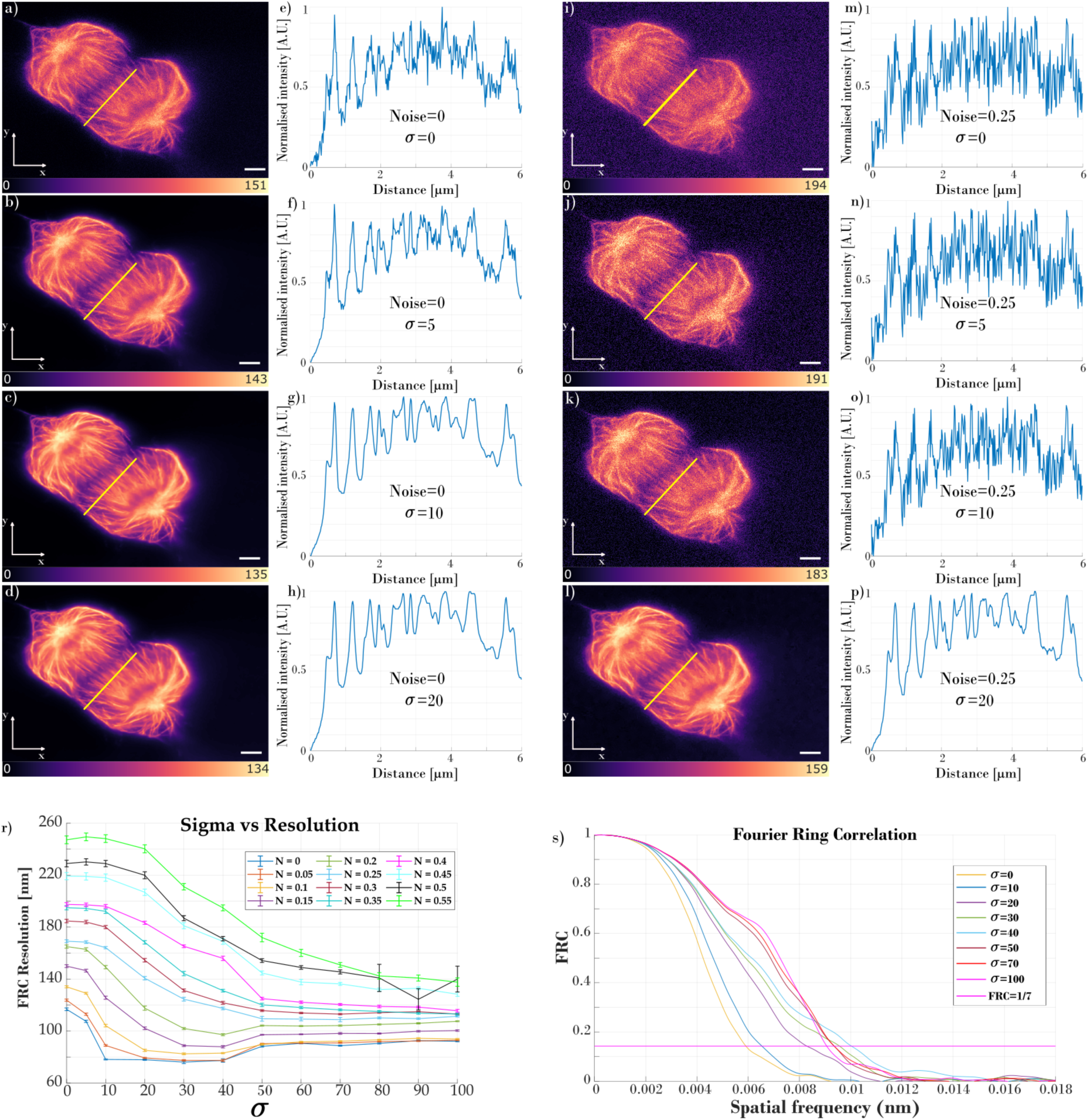
Analysis of the impact of the simulated noise and BM3D denoising on the effective image resolution. Random noise was simulated and added to images with different variance (from N = 0 to N= 0.55). The same images were denoised using BM3D algorithm with different value of sigma (from σ = 0 to σ =100). Images (a) – (d) show the analysis of acquired mitotic cell tubulin stained with Abberior STAR635P obtained using a 2D STED phase mask. (a) presents a raw image, (b) – (d) show the result of denoising with σ = 5, 10 and 20, respectively. Plots (e) – (h) show the intensity cross-sections marked with the yellow line in images (a) – (d), respectively. Images (i) – (l) show the analysis of the image (a) with the synthetic noise added with the magnitude N=0.25. Image (i) shows noisy data, images (j) – (l) demonstrate the effect of BM3D denoising with σ = 5, 10 and 20, respectively. Plots (m) – (p) show the intensity cross-sections marked with the yellow line in images (i) – (l), respectively. Plots in (r) present the analysis of the impact of different σ values for images with different noise level; (s) shows the FRC resolution obtained for the images denoised with different σ values for N=0.25. Scalebar size 2µm.

As shown in Fig. 6, BM3D filtering successfully retrieves image information covered in noise, but does not reject data that was originally present in the image, as the cross-section curve of the raw and denoised image are of the same shape. This feature can be seen as detail preservation and constitutes the main advantage of BM3D image denoising. According to the plots in Fig. 6(r) and 6(s), the σ value used in BM3D filtering that already enables satisfactory results is, depending on the noise level, between 5 and 10. Sigma ranging from 15 to 40 also provide generally favourable image denoising although it is recommended that this process is supervised through visual inspection (which was the case in preparing Fig. 6) to prevent loss of detail of fine structures under complicated real-life noise. Quantitative analysis of the FRC resolution with regard to the σ value shows that increasing noise significantly decreases the effective image FRC resolution, which can then be recovered using BM3D filtering. With added noise of variance N=0.2, the effective resolution decreases by 41%, from 117nm to 165nm; BM3D filtering with σ=20 returns the FRC resolution to 117nm. Similarly, analysing the intensity cross-sections through microtubules shown in Fig. 6(e)-(h) and Fig. 6(m)-(p), when the noise is raised to N=0.25, it is difficult to unambiguously state whether the imaged structure is information or noise; using BM3D filtering with σ=20, allows retrieval of the original information. In most of the cases, filtering with σ <10 is enough to successfully decrease the noise in the acquired image, therefore increasing its SNR.

For further analysis of the BM3D denoising performance, we used a real experimental image that has low SNR and high level of noise without resorting to additional synthetic noise generation. We chose an RPE-1 mitotic cell with its microtubules stained with the Abberior STAR635P. Figure 7 shows how important denoising is when looking at the images with insufficient SNR. When analysing the raw image (Fig. 7(a)), it is difficult to distinguish the noise from the object structure information and it is ambiguous whether the microtubules are visible or if it is the noise. For example, when looking at the intensity cross-section of the zoomed area in the raw image (Fig. 7(d)) the noise is at the same magnitude as the signal. However, when the images were processed using the BM3D algorithm, the microtubules are clearly visible both in the image and in the intensity cross-section. In this case filtering with σ=5 restores the fine image structures. The FRC analysis (Fig. 7(g)) proves that the resolution of the image increased from the initial 97nm to 87nm for filtering with σ=5 and to 76nm for σ=10. Moreover, Fig. 7 helps to highlight the fact that the combination of the FRC metric with visual inspection in some cases of real-life noise-corrupted fine structures is recommended. With a sufficiently high level of real-life noise, some very fine structures might be preserved better using σ=5 compared to σ=10, which provides a generally smoother image. Notably, denoising with both σ=5 and σ=10 enables significant improvements in image quality compared to the raw image – as it was confirmed by calculated FRC image resolution values.

**Fig. 7.**
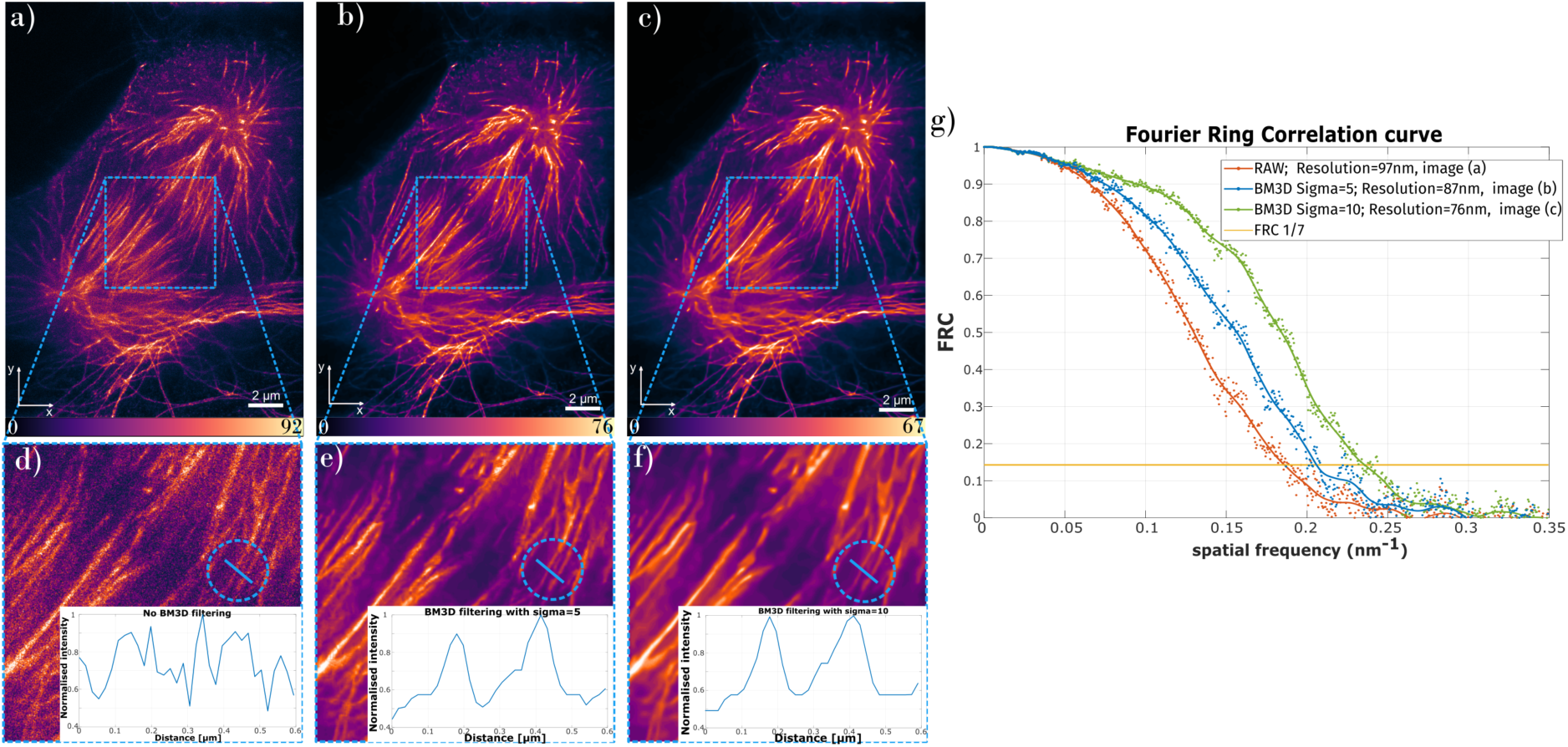
Resolution improvement on the noisy image using BM3D denoising algorithm. Mitotic cell tubulin was labelled with Abberior STAR635P and acquired using 2D STED phase mask. (a) raw, noisy image, (b) and (c) BM3D denoised image with σ=5 and σ=10, respectively. (d) – (e) show zoomed areas marked with blue, dashed rectangles in (a) – (c). The plots in insets in (d) – (f) are intensity cross-sections of the structures marked with blue line in images (d) – (e). Plots in (g) show FRC curves obtained for images (a) – (c).

Lastly, we used the BM3D algorithm to process the aberration corrected 3D STED images shown in Fig. 3, to improve the quality of the aberration corrected deep tissue image. Images from Fig. 3(c) were filtered using the BM3D denoising algorithm with σ=10. Comparison of the raw 3D STED and BM3D denoised 3D STED images can be seen in Fig. 8. The resulting BM3D processed images were much clearer both in the XY and XZ views, with the FRC resolution analysis of the unfiltered and filtered images indicating an improvement in FRC resolution from 182nm to 152nm in the BM3D filtered image for single slice. The average resolution for the whole axon volume increased from 204nm to 164nm. The FRC analysis showed that the noise has a strong impact on the final effective STED image resolution and that the noise filtering is beneficial to the quality of the imaging. Image acquisition is carried out by line-scanning of excitation and depletion beams across the sample – this fact is especially highlighted after BM3D denoising in Fig. 8(d).

**Fig. 8.**
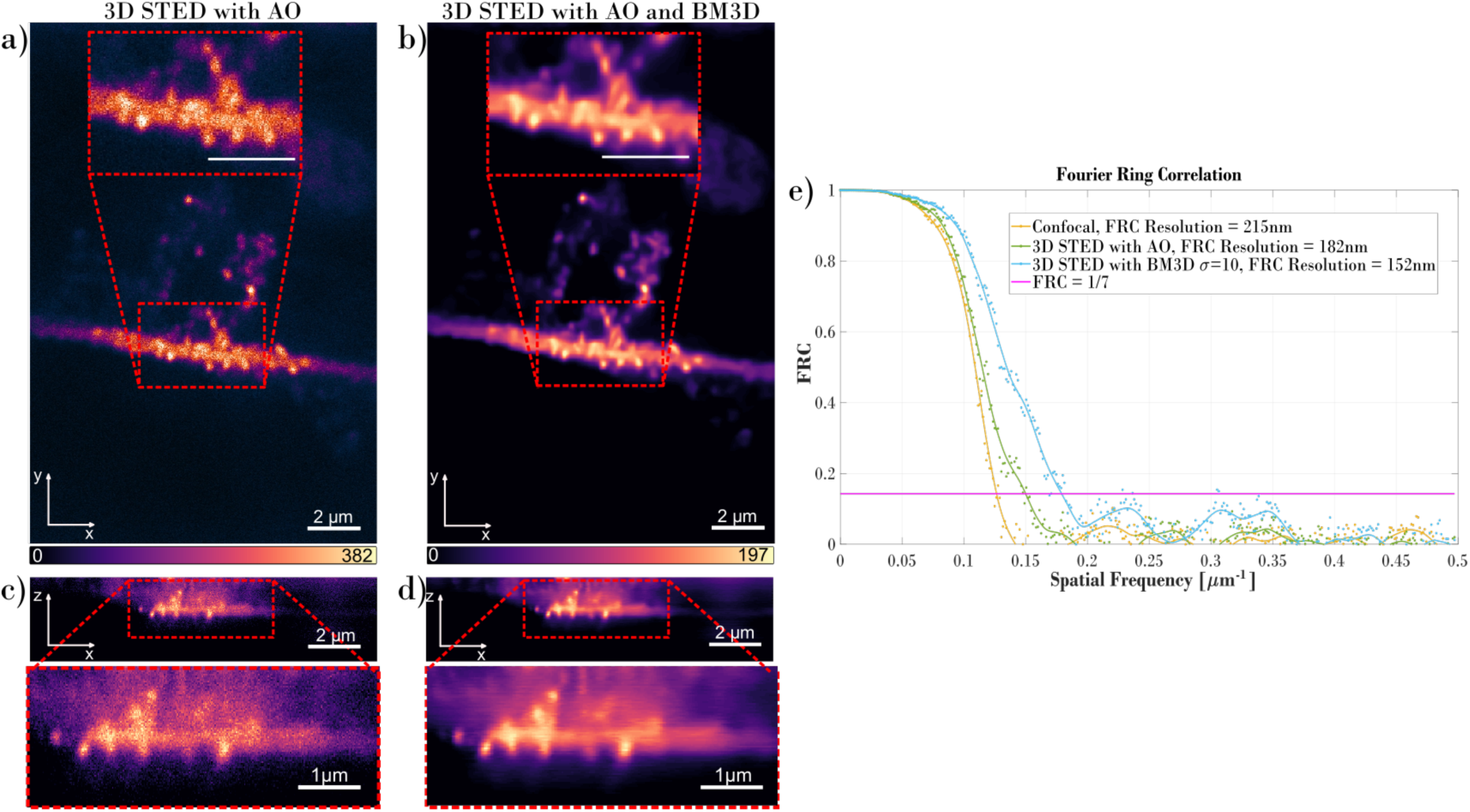
Effect of BM3D image denoising on the images shown in Fig. 3. (a) and (c) present the XY and XZ views, respectively, of an axon obtained with the 3D STED phase mask with aberrations corrected (as in Fig. 3 (c)). (b) and (d) are XY and XZ views, respectively, of the BM3D denoised images (a) and (c). Red dashed rectangles mark the area that is zoomed. Plots in (e) show the improvement of the effective FRC resolution when the acquired STED images are denoised using BM3D algorithm.

Fig. 9 presents the relationship between the sample depth and the FRC resolution of different imaging modes based on the exemplifying tissue studied in Fig. 3 and Fig. 8. The FRC resolution curves of aberration-corrected confocal (blue curve) and 3D AO STED (red curve) imaging modalities are alike and generally exhibit a slightly decreasing trend with increasing depth. An improvement can be seen for both imaging modalities with respect to the 3D STED without aberration correction (green curve). Additionally, the positive impact of the BM3D-based numerical enhancement on the 3D AO STED imaging resolution is clearly observable (orange and violet curves).

**Fig. 9.**
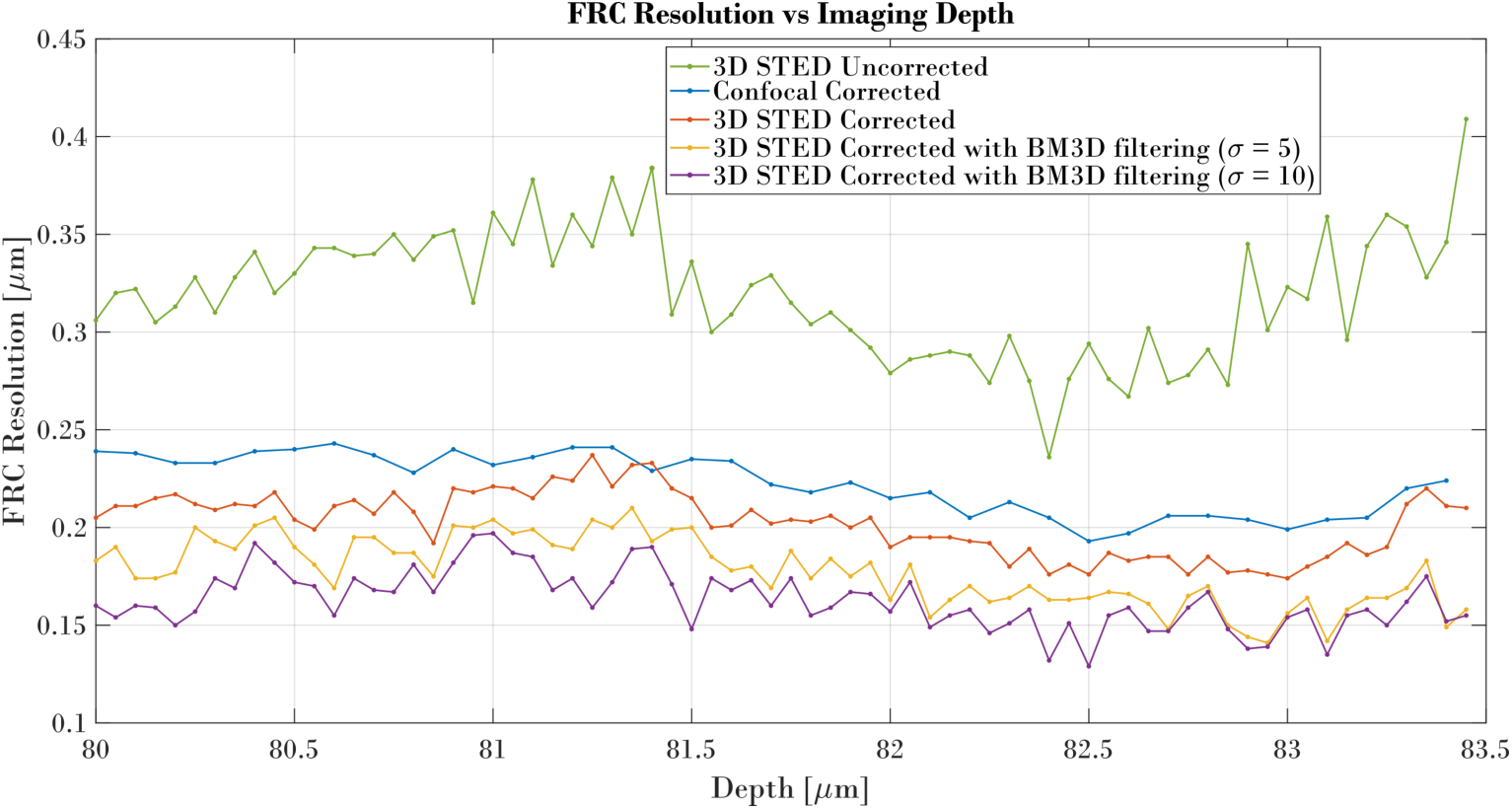
The relation between FRC resolution and the depth of the sample.

In Fig. 10 we show the comparison of the maximum intensity projection of the aberration corrected confocal and 3D STED stacks, with a depth color-coded colormap. Such comparison shows the qualitative improvement of the 3D STED images, as the colour diversity of the 3D STED image is stronger, and details sharper, compared with the confocal projection. Furthermore, we highlight that the numerical enhancement of the BM3D technique decreases noise-related granularity in the maximum intensity projection without jeopardizing the overall colour diversity and sharpness which can be seen as a very favourable outcome of image denoising.

**Fig. 10.**
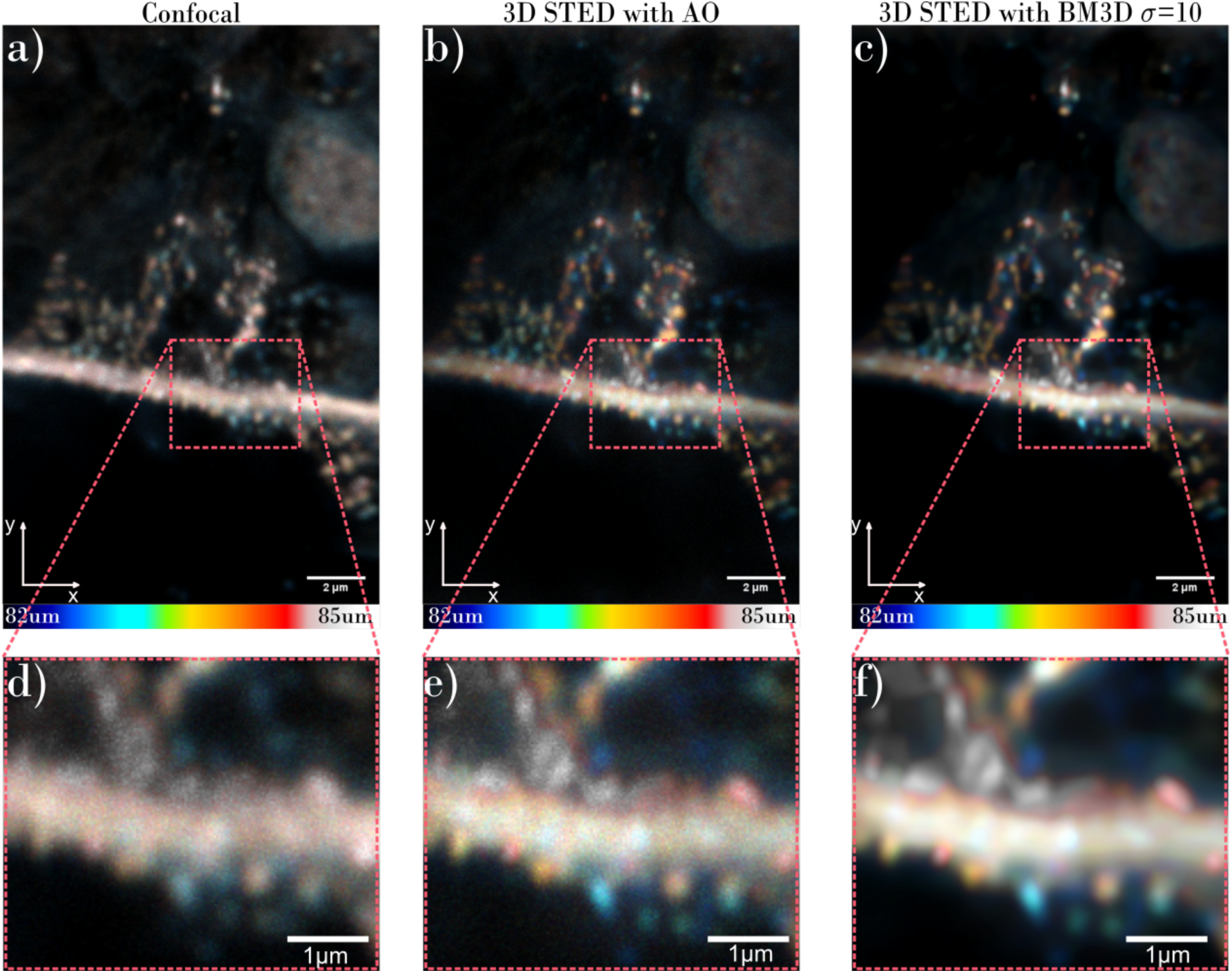
Comparison of the maximum intensity projections with the depth color-coded colormap of the (a) confocal, (b) proposed 3D AO STED and (c) 3D AO STED with BM3D images. Inserts (d), (e) and (f) show the zoomed areas marked with the red rectangles of (a), (b) and (c), respectively.

## Discussion

In this paper we demonstrated a custom-built STED microscope setup with dual adaptive optics that can be successfully used for super-resolved deep tissue imaging in thick samples. Thanks to the aberration correction of both excitation and depletion beams, we were able to image hiPSC-derived axons in DA neurons at a depth of 80µm, using a bottle beam for 3D STED imaging.

Fig. 3 shows the importance of correcting aberrations when imaging deep into an aberrating sample. Notably, confocal imaging with aberration correction offers higher quality imaging than STED imaging without aberration correction. It is worth emphasizing this fact, as super-resolution imaging of thick, optically dense specimen does not give satisfying results when the aberrations are not corrected. Using adaptive optics in the microscope we were able to correct aberrations introduced by the imaged sample and obtain the first super-resolved 3D STED images of an axon which grew under 80 microns of tissue. FWHM measurements indicate that the axial resolution of the corrected 3D STED images of the aberrating specimen was 2.5 times better than the confocal resolution (310nm for 3D AO STED and 795nm for confocal).

Thanks to the design of the microscope and the use of the resonant mirror we were able to successfully use the quantitative FRC metric for the evaluation of the effective lateral resolution of the microscope. STED images are acquired using high power laser light, which increases the noise level of the acquired image. Deep imaging through aberrating specimen adds another source of noise to the image, and as a result, the high spatial frequencies related to object structure (which correspond to the fine details of the image) are lost in the noise. Using the FRC metric for evaluation of the effective image resolution clearly showed that high laser-intensity-driven noise level can have a substantial effect on the decrease of final effective resolution of the acquired super-resolution image, as shown in Fig. 3.

We proposed employing the BM3D algorithm for the numerical denoising of the STED images in order to retrieve the object structure components that are covered by the optical imaging noise. In Fig. 6 we showed the analysis of the influence the BM3D filtering has on STED images with the controlled addition of synthetic noise. The results obtained in this carefully designed condition corroborated that BM3D can successfully retrieve the high spatial frequency details otherwise mixed with the noise without damaging the original object information (no blur is encountered). In Fig. 8 we presented the improvement of the image quality and effective FRC resolution when the raw image contains significant level of noise – BM3D filtering with σ=10 enabled over 20% of the FRC resolution increase. What is more, we showed that the combination of aberration correction and numerical enhancement (noise filtering) can successfully increase the quality, effective resolution and achievable depths of the super-resolution 3D STED imaging.

## Acknowledgements

This work was funded from the People Programme (Marie Curie Actions) of the European Union’s Seventh Framework Programme (FP7/2007-2013) under REA grant agreement no 608133. The microscope development was supported by the Wolfson Foundation and the Scottish Universities Physics Alliance (SUPA). P. Z. and M. T. would like to acknowledge the support of the National Science Center, Poland, under the project OPUS 13 (UMO-2017/25/B/ST7/02049), National Agency for Academic Exchange, and statutory funds of the Faculty of Mechatronics, Warsaw University of Technology. We thank Dr Iain Porter, Mr Michael Porter and Dr Lindsay Davidson for the preparation of the differentiated hiPSC-derived neurons.

