## Supplementary Material for "Numerically enhanced adaptive optics-based 3D STED microscopy for deep-tissue super-resolved imaging" (from another version of this preprint: version 1, 2019-05-29)

*Dual-SLM 3D STED microscope setup*

Our STED microscope schematic is presented in Fig. S1.

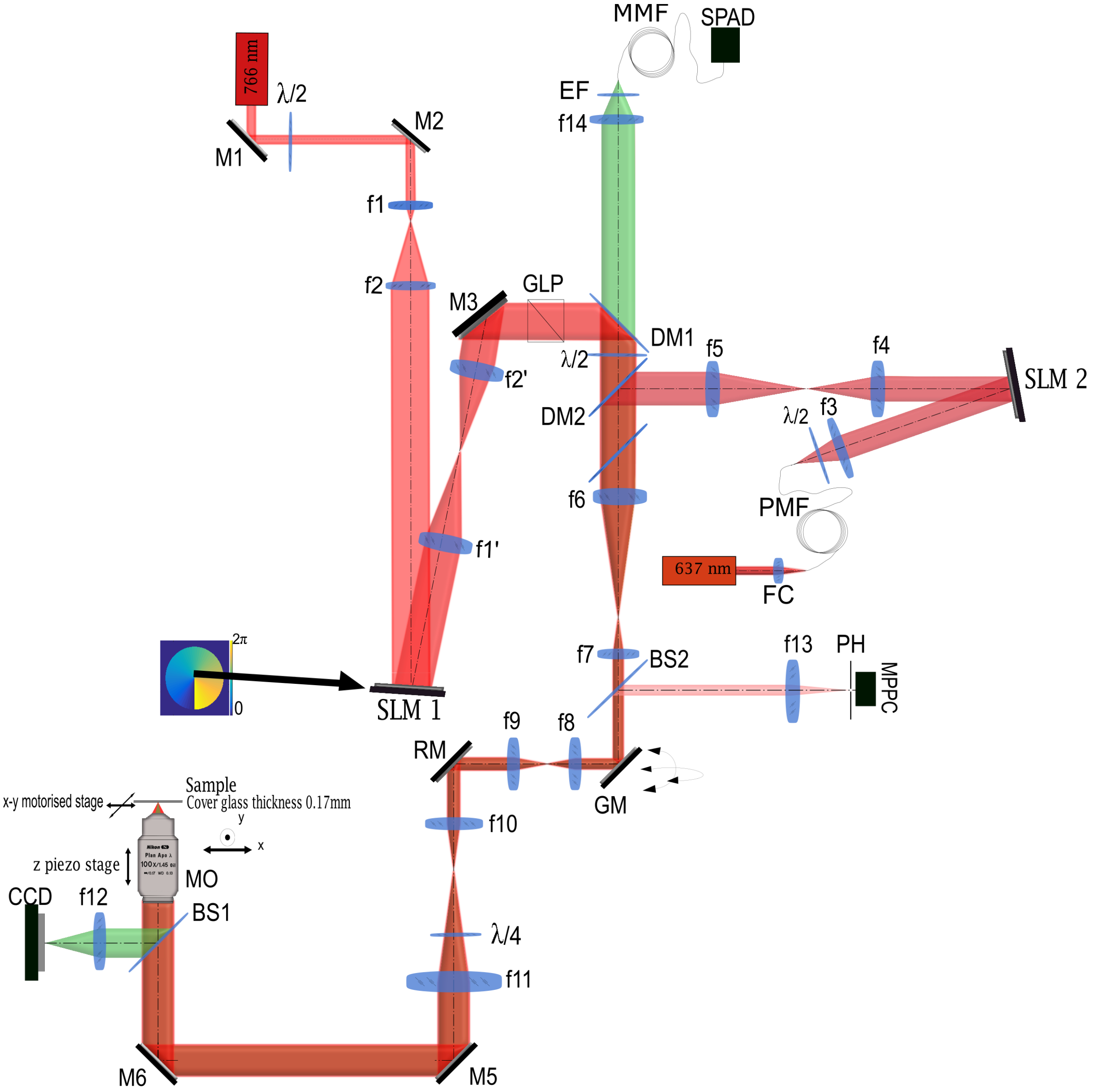


*Fig. S1. The optical setup of the custom built dual-SLM 3D STED microscope. 766nm – 766nm depletion laser, 637nm – 637nm excitation laser, BS1-BS2 – pellicle beamsplitters, CCD – CCD camera, DM1-DM2 dichroic mirrors, EF – emission bandpass filter, f1-f14 – lenses, FC – fibre coupling lens, GLP – glan laser polarizer, GM - pair of galvanometric mirrors, λ/2 – half-wave plate, λ/4 – quarter-wave plate, M1-M6 - steering mirrors, MMF – multi-mode fibre, MO - microscope objective, MPPD – multi pixel photon counter, PH – pinhole, PMF – polarization maintaining fibre, RM - resonant mirror, SLM 1 and SLM 2 - spatial light modulators, SPAD - single photon avalanche diode.*

**Depletion beam path.**

Depletion beam originates at the 766nm laser (PicoQuant VisIR 765 STED). It can be set manually to work at 80MHz, 40MHz, 20MHz, 10MHz, 5MHz, or 2.5MHz repetition rates and its pulse duration is approximately 0.5ns. Then it passes through the half-wave plate λ/2 (Thorlabs AQWP05M-600) and is expanded through lenses f1-f2 to match the size of the aperture of the liquid crystal on silicon spatial light modulator SLM1 (Hamamatsu X-10468-02). λ/2 rotates the depletion laser polarization so that it matches the orientation of the liquid crystal molecules of the SLM1. SLM1 is shaping the depletion beam into either 2D STED (Laguerre-Gaussian phase mask) or 3D STED (Top-Hat phase mask) beams. Pair of lenses f1'-f2' is imaging the SLM1 plane onto the dichroic mirror DM2 plane. Glan laser polarizer (GLP) is blocking any polarizations not corresponding to the orientation of the liquid crystals in the SLM1. Depletion beam is reflected into the main beam-path through the dichroic mirror DM1 (Semrock FF720-SDi01) and at the DM2 (Chroma ZT647rdc-UF3) it is superimposed with the excitation beam. λ/2 is used to rotate the polarization of the depletion beam. From the DM2 both depletion and excitation beams are co-propagating.

**Excitation beam path.**

Excitation beam is generated by the 637nm pulsed diode laser (PicoQuant LDH-P-C 640B). It operated at the same repetition rates that the depletion laser and has pulse width of 90ps. Excitation beam is coupled into the polarization maintaining single-mode fiber PMF using coupling lens FC. The output of the excitation beam has an elliptical shape and before coupling it has to be magnified over one axis using the anamorphic prism pair. This way the 75% of coupling efficiency can be obtained. λ/2 rotates the polarization of the depletion beam, lens f3 is used as collimator and such beam is then reflected from the SLM2 (Boulder Nonlinear Systems 512x512), which is used for correcting aberrations of the excitation beam. Lenses pair f4-f5 are imaging SLM2 plane onto the DM2 plane. DM2 is reflecting the excitation beam onto the main beam path and superimposes excitation beam with depletion beam.

**Main beam path**

Lenses f6 and f7 are imaging the DM2 plane onto the plane located between closely aligned pair of galvanometric mirrors GM (ScanLab Dynaxis XS) and lenses f8 and f9 image the GM plane onto the 16kHz resonant mirror RM (Electro-Optical Products Corp. SC-30) plane. Combination of GM and RM is used for scanning the sample – RM scans fast axis, GM scans slow axis. Lenses f10 and f11 are imaging RM plane onto the back focal plane of the microscope objective MO (Nikon CFI Plan Apo Lambda 100X Oil NA 1.45). MO is placed on the focusing piezo stage (Piezoconcept HS1.70) while sample is mounted on the motorized XY stage (ASI Imaging ASI S31121010FT).

**Imaging beam paths.**

There are 3 imaging modes: brightfield transmission mode, quasi-widefield reflection mode and confocal fluorescent mode.

For imaging using a brightfield mode, sample is illuminated from top using white light diode. Transmitted light is directed onto the CCD camera using 92:8 pellicle beamsplitter BS1 and focused using tube lens f12.

Reflected light, for imaging reflective objects such as gold beads, is de-scanned using RM and GM mirrors and then reflected using 92:8 pellicle beams splitter BS2. Then, lens f13 is focusing the reflected light through the widefield pinhole. Multi Pixel Photon Counter MPPC (Hamamatsu C13366).

Fluorescent light is de-scanned using the RM and GM mirrors, separated from the excitation beam by BM2 and the separated from the depletion beam by DM1. Lens f14 is focusing the fluorescent signal into the multi-mode fiber MMF, which acts as a confocal pinhole. The size of the MMF is set to 0.8 Airy units. Bandpass emission filter EF (Semrock 676/37) is rejecting any remaining non-fluorescent signal.
